## Supplemental material for "Characterization of a new lacrimal gland cell line in 2D and 3D cell culture models"

Additional material for

Sophie Gleixner *et al.*

**This PDF file includes:**

Additional Figures 1 to 4
Additional Table 1

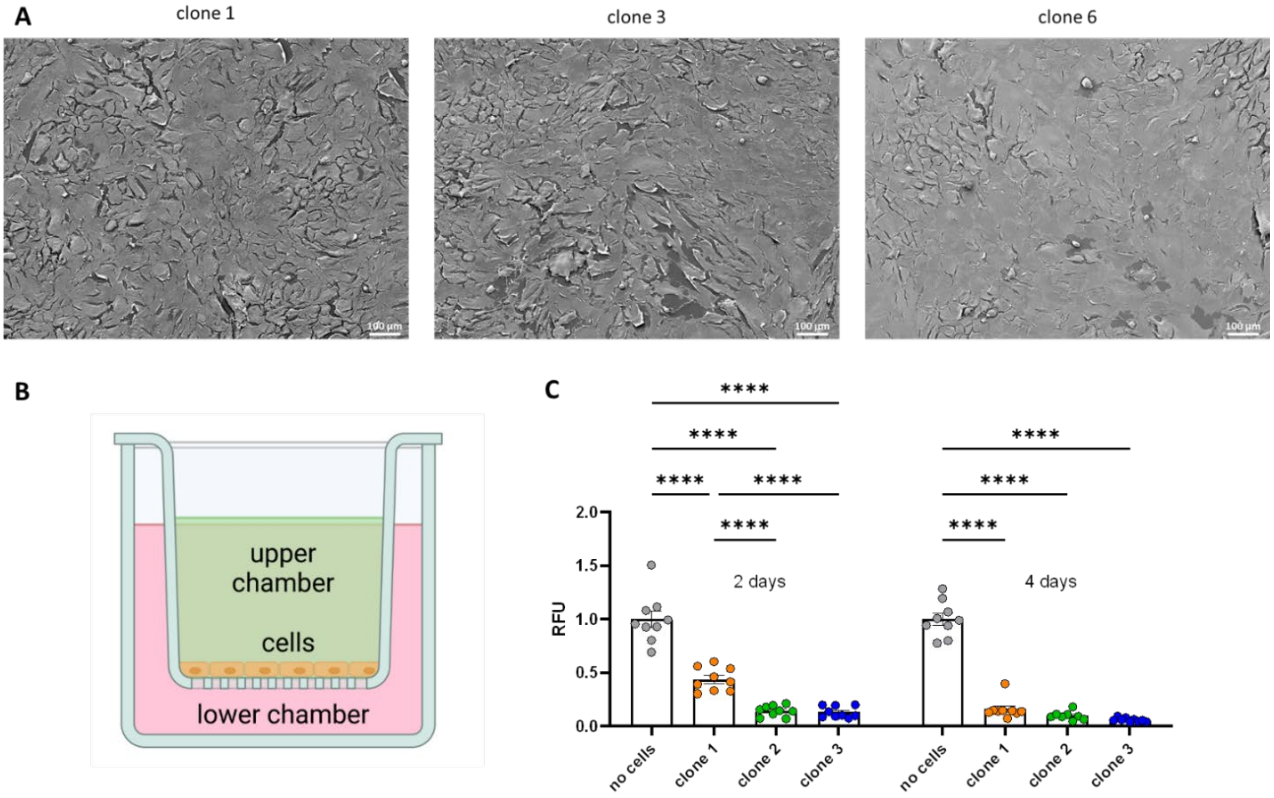


**Additional Figure 1.**

Validation of the epithelial character of three immortalized human lacrimal gland epithelial cell clones**. A**) Scanning electron microscopy of the surface of clone 1, 3 and 6 after cells reached confluence. **B**) Cartoon of FITC-dextran permeability assay setup. The figure is partly created with BioRender.com. **C**) Relative fluorescence units (RFU) measured after 30 minutes in the lower chamber of wells with cell culture inserts containing cells of clone 1, 3 and 6 after 2 or 4 days of growth. Data are means ± SEM [*$p < 0.05$, **$p < 0.01$, ***$p < 0.001$, and ****$p < 0.0001$, Two-way ANOVA, N = 3].

| gene | protein |
|---|---|
| AQP5 | Aquaporin 5 |
| CST6 | Cystatin 6 |
| FOXC1 | Forkhead Box C1 |
| MYL9 | Myosin Light Chain 9 |
| PAX6 | Paired Box Gene 6 |
| CSTB | Cystatin B |
| VWF | Von Willebrand Factor |
| CD14 | Monocyte Differentiation Antigen CD14 |
| CD34 | Sialomucine |
| ACTA2 | Smooth Muscle Actin Beta 2 |
| HAS2 | Hyaluronan Synthase 2 |
| FN1 | Fibronectin 1 |
| DES | Desmin |
| PALLD | Palladin |
| VIM | Vimentin |
| POSTN | Periostin |
| LUM | Lumican |
| PDGFRB | Platelet Derived Growth Factor Receptor Beta |
| MFAP5 | Microfibril Associated Protein 5 |
| COL1A2 | Collagen Type I Alpha 2 Chain |
| COL6A2 | Collagen Type VI Alpha 2 Chain |

**Additional Table 1.**

List of marker genes for lacrimal gland epithelial cells, endothelial cells, mesenchymal stem cells, myoepithelial cells and fibroblasts.

A

| gene | protein |
|---|---|
| CST6 | Cystatin 6 |
| FOXC1 | Forkhead Box C1 |
| MYL9 | Myosin Light Chain 9 |
| PAX6 | Paired Box Gene 6 |
| CSTB | Cystatin B |
| ACTA2 | Smooth Muscle Actin Beta 2 |
| C3 | Complement C3 |
| KRT5 | Keratin 5 |
| TNFRSF19 | TNF Receptor Superfamily Member 19 |
| AQP5 | Aquaporin 5 |
| CHRM3 | Cholinergic Receptor Muscarinic 3 |
| ADRB1 | Adrenoceptor Beta 1 |
| HTRA4 | HtrA Serine Peptidase 4 |
| MYH11 | Myosin Heavy Chain 11 |
| LCN2 | Lipocalin 2 |
| VIPR1 | Vasoactive Intestinal Peptide Receptor 1 |
| PIGR | Polymeric Immunoglobulin Receptor |
| SFTPD | Surfactant Protein D |
| LTF | Lactoferrin |
| LGR5 | Leucine Rich Repeat Containing G Protein-Coupled Receptor 5 |
| SERPINB2 | Serpin Family B Member 2 |

B

| gene | protein | size [bp] |
|---|---|---|
| ACTA2 | Smooth Muscle Actin Beta 2 | 88 |
| ACTB | Beta Actin | 298 |
| AQP5 | Aquaporin 5 | 232 |
| CST6 | Cystatin 6 | 255 |
| CSTB | Cystatin B | 243 |
| FOXC1 | Forkhead Box C1 | 82 |
| MYL9 | Myosin Light Chain 9 | 334 |
| PAX6 | Paired Box Gene 6 | 500 |

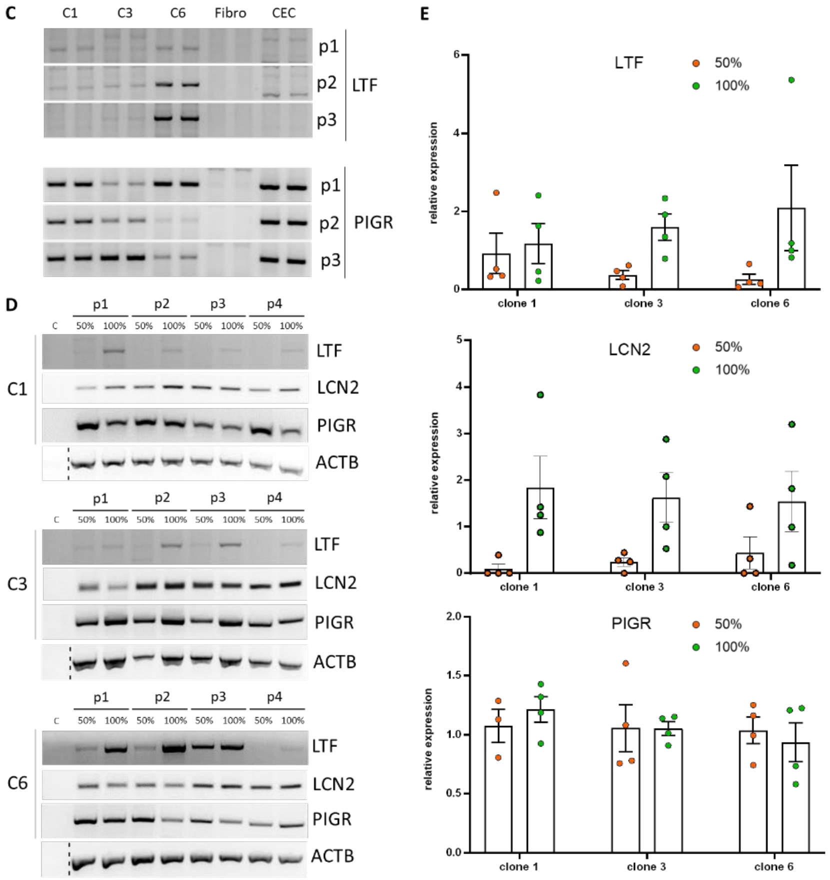


**Additional Figure 2.**

Comparison of expression of different lacrimal gland marker genes. **A**) List of lacrimal gland marker genes. **B**) List of the marker genes tested in Fig. 3B with gene names, names of the respective protein product and the expected amplicon size. **C**) Agarose gels with the amplification products of RT-PCR for the tear fluid proteins lactoferrin (LTF) and Polymeric Immunoglobulin Receptor (PIGR) at three different passages (p1, p2, p3) for clone 1 (C1), 3 (C3), 6 (C6), human fibroblasts (Fibro) and human corneal epithelial cells (CEC). **D**) Agarose gels with the amplification products of RT-PCR for LTF, LCN2, PIGR and actin beta (ACTB) at 50 % and 100 % confluence at four different passages (p1-p4) for clone 1, 3 and 6 and a negative control (C). **E**) Semiquantitative comparison of LTF, LCN2 and PIGR expression level in clone 1, 3 and 6 at 50 % and 100 % confluence. Data are means ± SEM [Two-way ANOVA, N = 4].

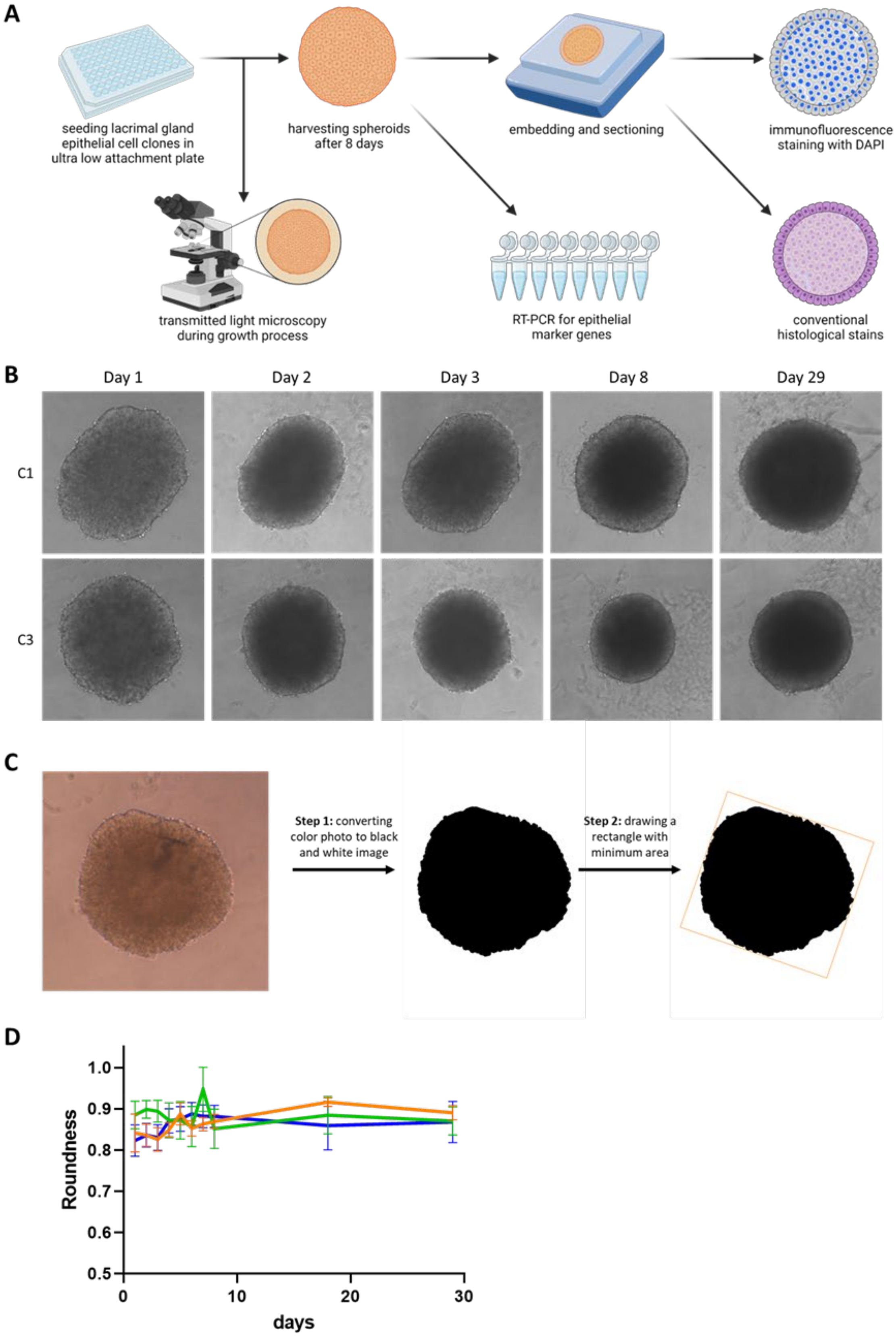
A
seeding lacrimal gland epithelial cell clones in ultra low attachment plate
harvesting spheroids after 8 days
embedding and sectioning
immunofluorescence staining with DAPI
transmitted light microscopy during growth process
RT-PCR for epithelial marker genes
conventional histological stains
B
Day 1
Day 2
Day 3
Day 8
Day 29
C1
C3
C
Step 1: converting color photo to black and white image
Step 2: drawing a rectangle with minimum area
D
Roundness
1.0
0.9
0.8
0.7
0.6
0.5
0
10
20
30
days

**Additional Figure 3.**

Analysis of spheroidal growth. **A**) Cartoon illustrating the workflow from the lacrimal gland epithelial cells clones to 3D spheroids and their analysis. The figure was partly created with BioRender.com. **B**) Transmitted light microscopy during the 29 days culture period of 3D grown cells of clone 1 (C1) and 3 (C3). **C**) To determine the circularity of the spheroids, in the first step the microscopic images were converted to black and white images for representing each spheroid as a black object on white background. In the second step, these images were analyzed using an in-house Python script for measuring the size of objects in an image with OpenCV. This script draws a rectangle with the minimum area around the black object and calculates the edge lengths of the rectangle. For exactly circular objects, both edges of the rectangle have the same length, while for elliptical objects the edge lengths differ. **D)** Roundness of the spheroids of clone 1 (orange), 3 (green) and 6 (blue) over the course of the 29 days culture period. Data are means ± SEM. [Day 1-8 N = 24, day 18 N= 9, and 29 N = 9].

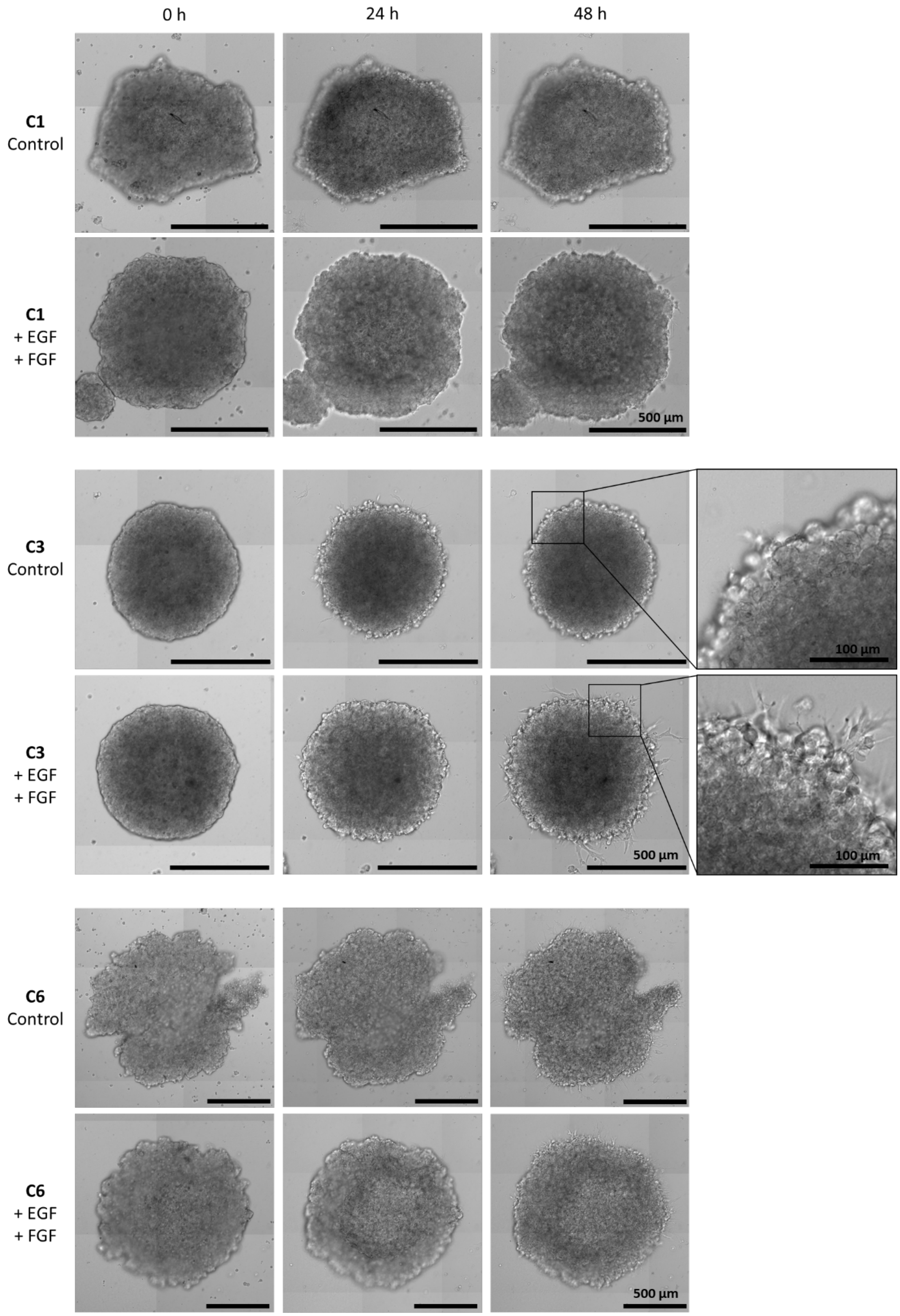
0 h
24 h
48 h
C1
Control
C1
+ EGF
+ FGF
500 µm
C3
Control
100 µm
C3
+ EGF
+ FGF
500 µm
100 µm
C6
Control
C6
+ EGF
+ FGF
500 µm

**Additional Figure 4.**

Spheroids in extracellular matrix with and without growth factors (epidermal growth factor (EGF) and fibroblast growth factor 10 (FGF10)) added to the medium form buds after 0, 24 and 48 hours.
